# Current geography masks dynamic history of gene flow during speciation in northern Australian birds

**DOI:** 10.1101/178475

**Authors:** Joshua V. Peñalba, Leo Joseph, Craig Moritz

**Author notes:** Corresponding author: Joshua Penalba.

## Abstract

During early stages of speciation, genome divergence is greatly influenced by gene flow. As populations diverge, geography can allow or restrict gene flow in the form of barriers. Current geography, e.g. whether sister species are allopatric or parapatric, is often used to predict the potential for gene flow during the divergence process. We test the validity of this assumption in eight meliphagoid bird species codistributed across four regions. These regions are separated by known biogeographic barriers within and between northern Australia and Papua New Guinea. We find that bird populations across the same barrier have a range of divergence levels and probability of gene flow regardless of range connectivity. Geographic distance and maximum range connectivity over time can better predict divergence and probability of gene flow than whether populations are currently allopatric or parapatric. We also find support for a nonlinear decrease of the probability of gene flow during the divergence process. This implies that although gene flow influences divergence early in speciation, other factors associated with higher divergence restrict gene flow later in speciation. Current geography may then mislead inferences regarding potential for gene flow during speciation under a complex and dynamic history of geographic and reproductive isolation.

## Background

Gene flow, selection, and genetic drift shape divergence during speciation, while geography sets the stage on which these forces act [1–3]. The geographic mode of speciation is defined by the extent of spatial isolation during early stages of divergence. Population pairs can have disjoint (allopatric), completely overlapping (sympatric), or separate yet partially adjoining (parapatric) ranges [4]. This geographic context predicts potential gene flow between populations. Variation in levels of gene flow affects genetic differentiation, in turn affecting the strength of selection and drift that is necessary to drive population divergence. Alternatively, under a purely population genetic framework the geographic mode of speciation is defined by levels of gene flow; “allopatry” when the proportion of the population which are migrants (*m*) equals zero, “sympatry” when *m* = 0.5, and “parapatry” when 0 < *m* < 0.5 per generation [5].

Although geographic and genetic definitions are often assumed to correspond, this is not always the case in nature [6]. Current range distributions between sister species are often used to infer their geographic mode of speciation, although the dynamic nature of species ranges through evolutionary time can lead to unreliable predictions of gene flow [7]. Range fluctuations during the Pleistocene’s climatic cycling resulted in multiple periods of connectivity and discontinuity before resulting in the distribution we see today [8]. In order to understand how gene flow has influenced divergence during speciation, we must first understand how the geographic history could have shaped the potential for gene flow through time. The discrepancy between the spatial and population genetic definitions of the geographic mode begs the question: does current geography adequately predict realized gene flow and, consequently, divergence during early stages of speciation?

Although geographic connectivity would allow for gene flow early in speciation, later in the proces geographic connectivity can be insufficient for gene flow as populations diverge and become reproductively isolated. Populations further along in the speciation process would have reduced realized gene flow due to intrinsic incompatibilities or extrinsic selection [9]. Especially in birds, prezygotic isolation from sexual selection on song or plumage could influence gene flow in later stages of speciation. There has been growing support for the “snowball” model of accumulating incompatibility loci, initially proposed by Orr [10]. Qualitatively, this model and those derived from it specify a nonlinear accumulation of incompatibility loci resulting in a short speciation duration [11]. The “snowballing” has also been modeled under scenarios with moderate to no gene flow (parapatry to allopatry) meaning different geographic modes may yield a similar short duration of speciation, varying only in how long it takes for speciation to initiate [12–16]. This rapid accumulation of isolation has also been proposed under models of divergent selection in speciation-with-gene flow and under neutral models [17–20]. Though the underlying assumptions of these theories may differ, the trajectory converges to a rapid transition, ‘ snowballing’, or ‘ tipping point’ during speciation (simply referred to ‘ snowballing’ from this point forward). There is increasing empirical support for this pattern from studies of individual systems [21–23] and a taxonomically broad meta-analysis [24]. More broad, comparative studies across multiple systems would help elucidate this trajectory to speciation.

Further studies of population divergence in various geographic contexts could further clarify the role of gene flow, selection and genome architecture in influencing the landscape of divergence [9,20,25,26]. During speciation, local genomic variation in mutation and recombination rates influence the rate at which regions diverge [25,27,28]. The geographic mode of speciation could influence this landscape of divergence. Theory predicts that populations diverging in parapatry should have a skewed distribution of divergence across the genome with a few loci resistant to gene flow due to selection while the rest are free to move between populations [29]. Populations diverging in allopatry, on the other hand, are predicted to lack this skew as drift would be the predominant force influencing divergence [30,31]. This landscape should also change as populations move further along the speciation continuum [9,27].

Here we investigate multiple bird species with populations codistributed in the same region with known biogegraphic barriers. Empirical studies of gene flow during divergence in relation to geography often survey closely related populations and species across different geographic regions with varying biogeographic histories and selection pressures [30,31]. However, to understand how shifting geographic ranges can erode the correspondence between current geography and realized gene flow, it is more relevant to compare a set of taxa across common geography. Our study region comprises part of the monsoonal tropics of northern Australia and southern Papua New Guinea containing congruent biogeographic barriers for many taxa including birds [32–34]. Sea level rise since the last glacial maximum has formed a barrier between northern Australia and Papua New Guinea [35]. Meanwhile, the aridification of mainland Australia has resulted in multiple semipermeable terrestrial barriers with parapatrically or allopatrically distributed populations. Multiple studies have shown congruent phenotypic and genetic breaks for various plant and animal species in this region [34,36–39]. Here we use one gerygone and seven honeyeater species co-distributed in four focal regions: Papua New Guinea (PNG), Northern Territory (NT), Cape York Peninsula (CYP) and eastern Queensland (QLD) that are separated by well-known barriers (figure 1). These species were chosen as they have already been shown to have variation in divergence levels across known barriers between CYP and QLD [40] and have varying degrees of range connectivity [41] setting the stage to compare divergence in different biogeographical contexts. With this system we ask (1) how current geography, or potential for gene flow, predicts realized gene flow and genome divergence during early stages of speciation, (2) how genome divergence influences realized gene flow in later stages of speciation, (3) and how divergence varies across different loci during the speciation process. Focusing on the codistributed ranges of these species will allow us to understand how the genome differentiates during speciation under different geographic contexts within and between closely related systems.

**Figure 1.**
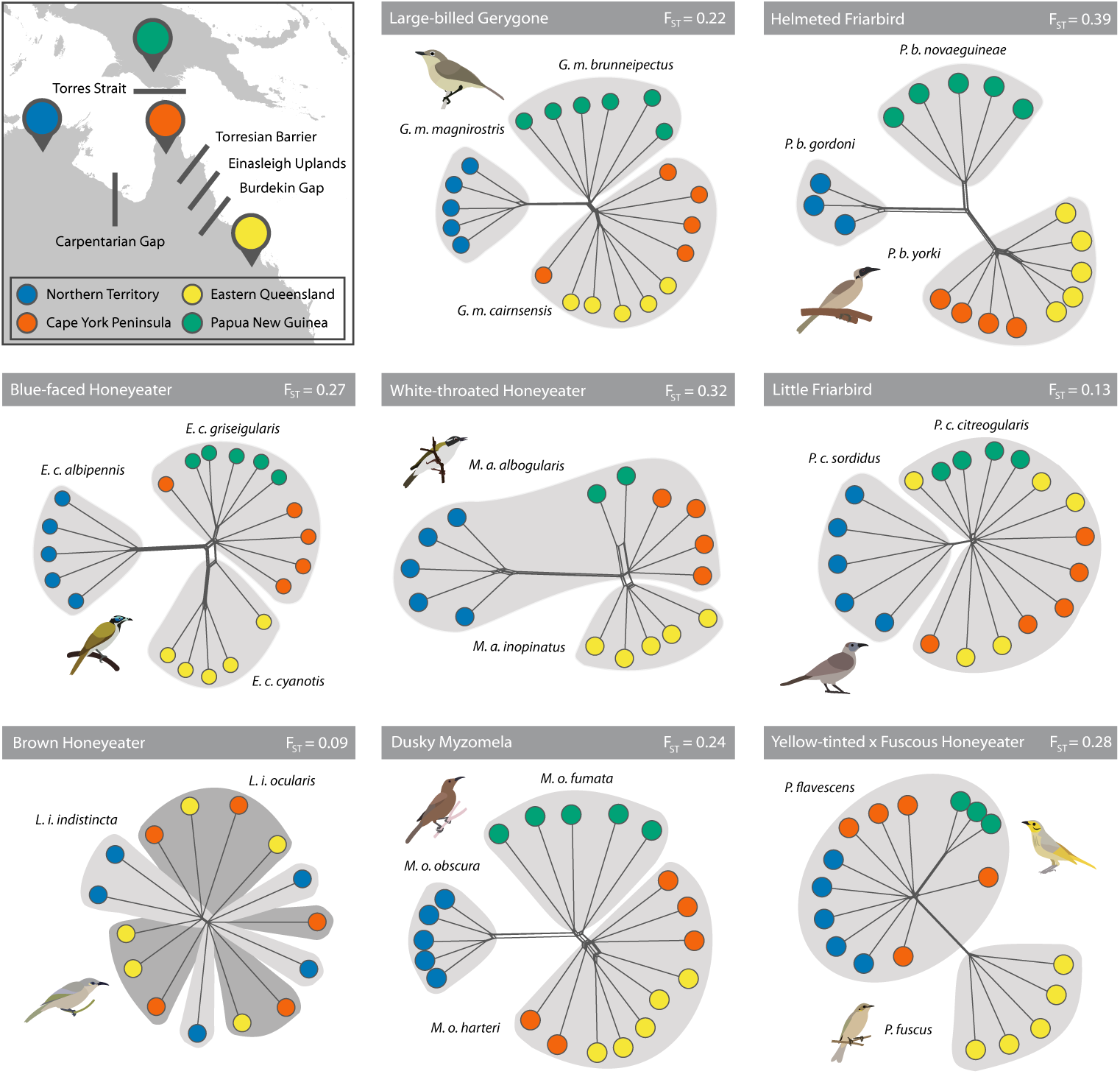
Geographic regions that were sampled for this study with corresponding known biogeographical barriers. Sample networks were generated from a distance matrix estimated from the ddRAD genotype likelihoods. Within each system, the species and subspecies designation has been highlighted. F_ST_ represents global measures across all four populations.

## Methods

### Sampling

We sampled eight species with populations that occupy the four regions of interest (figure 1). For each population, we sampled three to six individuals for a total of 157 individuals (15-20 individuals per species; electronic supplementary material, table S1). Samples were chosen from locations farther away from known contact zones to avoid recent hybrids [42]. Brown honeyeater and white-throated honeyeater had insufficient sampling for PNG so only the Australian populations were used in the analyses. We extracted the DNA from all individuals using a standard salting-out procedure.

### Sequencing

We sanger sequenced NADH dehydrogenase-2 (ND2) using the primers L5204 (5’ TAACTAAGCTATCGGGCGCAT 3’) and H6312 (5’CTTATTTAAGGCTTTGAAGGCC 3’) for measures of mitochondrial divergence and structure [43]. For the nuclear loci, we used a slightly modified version of the ddRADseq protocol as described by [44]. In brief, we digested the DNA using the restriction enzymes *PstI* and *EcoRI* and size-selected 345 - 407bp. Approximately ten indexed individuals in ten pools were sequenced on a NextSeq500 for 150bp, single-read, mid-output and the rest were sequenced on another NextSeq500 lane with similar specification except with high-output. A more detailed description of lab methods is available in the electronic supplementary material.

### Data processing and analyses

For each species we generated a reference set of RAD loci via the pyrad pipeline [45]. We used individuals from a different species to serve as an outgroup to polarize the SNPs in downstream analyses. Only RAD loci which have associated outgroup sequences were retained for the reference (electronic supplementary material, table S2). The resulting set of sequences were used as a reference for further processing. Individual reads were then mapped onto this reference set using Bowtie2 (v. 2.2.2)[46]. In order to recover larger numbers of loci and avoid biases from filtering for loci with higher coverage, the mapped reads were further processed using ngsTools and ANGSD to incorporate genotype likelihood information for the various population genetic measures (v. 0.911; http://www.popgen.dk/angsd) [47].

### Population structure

It has been demonstrated that more loci, even of lower coverage, yields more accurate estimates of population genetic statistics compared to fewer loci of higher coverage when using genotype likelihoods [48].We used ANGSD to further filter for SNPs to be used for downstream analyses (electronic supplementary material, table S3; see electronic supplementary material for detailed ANGSD commands). We used a minimum coverage cut-off of 2X and a maximum cut-off of 40X per individual to optimize the number of loci to be used while reducing the likelihood of recovering paralogous loci. The maximum coverage cut-off was determined by plotting a histogram of the average coverage per RAD locus and finding the upper threshold where most loci fell under. To determine population structure, we randomly chose a single SNP per RAD locus and reran ANGSD only for that set. We then used used ngsDist [49] to generate a distance matrix which we used for a principal coordinates analysis (PCoA) using ‘ cmdscale’ from base R (v3.2.2) and a population network using SplitsTree (figure 1, electronic supplementary material figure S1) [50].

### Population divergence statistics

To calculate the various population divergence statistics (F_ST_, D_XY_, and D_A_) we used the software within the ngsTools package. These statistics were used as they provide both relative and absolute measures of genetic divergence and can be compared to previous studies [24,30,31].

We used all SNPs within each RAD locus for all per locus and global measures. To calculate pairwise F_ST_, we used realSFS on the ANGSD genotype likelihoods to estimate an unfolded 2D site frequency spectrum (SFS) per population pair and used the SFS to derive the per locus F_ST_ estimate [51,52]. We used the same outgroups as the *pyrad* filtering steps for the unfolded SFS. To calculate global F_ST_, we used the estimate of allele frequencies within each population and all populations pooled together F_ST_ = H_T_ - H_S_ / H_T_ where H_T_ is the heterozygosity of all populations pooled and H_S_ = ΣH_e_ / k and H_e_ = 1 – Σ(p^2^ + (1-p)^2^)/m where k is the number of populations, m is the number of loci, and p is the allele frequency [53]. To calculate D_XY_, we used the estimate of allele frequencies which incorporated the genotype likelihood from ANGSD [54]. To calculate D_A_, we used the π estimates from ANGSD for each population and the equation D_A_ = D_XY_ - (π_X_ + π_Y_)/2. Lastly, we used the package ape v. 4.1 to calculate the ND2 genetic p-distance under the Jukes-Cantor model [55].

### Estimating the likelihood of migration

We used an approximate Bayesian computation (ABC) model selection to estimate the likelihood of migration (i.e. how strong the evidence is for some gene flow during the divergence process) between each population pair. The ABC analyses and models followed that of Roux et al. but using the unfolded 2DSFS as a summary statistic instead of the various population genetic statistics used in their study [24]. We tested the models of isolation-with-migration (IM), isolation-with-migration + heterogeneous Ne (IMhetN), isolation-with-migration + heterogeneous migration (IMhetM), isolation-with-migration + heterogeneous Ne + heterogeneous migration (IMhetNhetM), strict isolation (SI), and strict isolation + heterogeneous Ne (SIhetN). Heterogeneous Ne is modeled to reflect variation in recombination rate throughout the genome and heterogeneous migration is to reflect variation in gene flow across loci between hybridizing populations. Models of SI preceding IM (secondary contact) and IM preceding SI were excluded as they are typically difficult to distinguish from IM models [24]. We then used the R package abc v. 2.1 and calculated the likelihood of each model using a neural network with 50 trained and 6 hidden networks [56]. We ran the abc analyses five times and used the average model support of the replicates for further analyses. As in Roux et al., we used the sum of the support for models containing a migration parameter as our probability for migration [24]. Additional details of the analyses can be found in the supplementary material.

To infer the relationship between divergence and realized gene flow, we correlated F_ST_ to the probability for migration between each population pair. To negate possible circularity where model support may be determined by F_ST_ itself, we calculated the F_ST_ of simulations under the SI model and reran the ABC model selection on a range of F_ST_ values. If the ABC model selection consistently recovers a low probability of migration, regardless of F_ST_, high support for IM for populations with lower F_ST_ in the empirical data would more likely be a biological phenomenon rather than a model selection artifact.

### Speciation model fitting

We fit our divergence versus probability of migration distribution to test support for various theoretical trajectories of parapatric speciation presented by Yamaguchi and Iwasa [15]. The models include (1) ‘ threshold’: where full incompatibility is reached after a certain divergence level, (2) ‘ constant’ rate of divergence increase, (3) ‘ accelerated’ where increase in divergence is small until a certain divergence threshold is reached and increase is accelerated, (4) ‘ decelerated’ increase where divergence accumulates quickly but slows down as it approaches full incompatibility, and (5) ‘ sigmoid’ where the rate of increase starts slow but accelerates after a certain threshold before decelerating again prior to reaching complete incompatibility suggesting a snowballing during the speciation process (electronic supplementary material, table S4, figure S2). Our global F_ST_ parallels the ‘ incompatibility genetic distance’ of Yamaguchi and Iwasa [15]. The inverse of our probability of migration parallels their ‘ incompatibility’ where no support for migration (*P*(*m*) = 0)) is equivalent to one or full support for incompatibility (*I*(z) = 1). Identical to F_ST_ and probability of migration, the measures Yamaguchi and Iwasa used are bounded by zero and one. We used the resulting probability of migration as the summary statistic and simulated 500k distributions under each speciation trajectory model and used ABC and model rejection to estimate model support for our data. To test robustness of our ABC model inference, we used 200 simulated datasets under each model to see if the correct model is recovered (electronic supplementary material, table S5). Details regarding this method can be found in the supplementary material.

### Species distribution modeling

To infer how geographic distributions have changed through time we estimated geographic connectivity over space through time using species distribution modeling and least cost path analyses. We used vouchered specimen data from the Atlas of Living Australia (http://www.ala.org.au) for occurrence points and environmental variables from WorldClim v. 1.4 (Community Climate System Model 4; http://worldclim.org/paleo-climate1) to predict species ranges under past climates. To model species distributions in R v. 3.2.2, we followed guidelines described in [57]. We used the R package dismo v. 1.1-1 for the maximum entropy (MAXENT) analyses to predict species ranges [57]. We ran MAXENT using environmental layers from the present, mid-Holocene and the LGM (electronic supplementary material, table S6). Lastly, we selected a single coordinate (midpoint of the range) for each of the four populations and calculated the geographic distance between those points to account for isolation-by-distance. Between population pairs, we also calculated the least cost path using the R package gdistance (v. 1.1-9) based on modeled suitability [58] in the current range, mid-Holocene, and LGM predictions. For each population pair, we chose the minimum resistance path between all three time points to quantify the highest opportunity for migration through time. Habitat resistance values can be found in the electronic supplementary material table S7 and species distribution predictions can be found in figure S3.

## Results

### Population structure and divergence

Population structure varied between each taxon-pair though most had some form of clustering across the four populations (figure 1). Of the eight species, five showed distinct clusters for all four populations, one showed only three distinct clusters, one showed only two distinct clusters, and the last showed no population clusters (F_ST_ 0.09 - 0.39). The degrees of clustering and the population relationships are fairly variable, as shown by a PCoA and population network of the genetic distances (electronic supplementary material, figure S1). Currently allopatric populations are not more likely to be separate clusters, exemplified by samples from the PNG population often being closer to CYP and QLD. Similarly, geographically parapatric populations are not necessarily mixed as as shown by NT often being the most diverged and consistently separate cluster even for species with currently continuous geographic ranges to CYP and QLD. Mitochondrial haplotype networks generally corroborated nuclear SNP structure. Measures of genetic differentiation and divergence also varied. Autosomal and Z chromosome divergences were consistently correlated with all divergence measures (electronic supplementary material, figure S4, tables S8 and S9). Relative divergence (F_ST_) and gene flow scales with mitochondrial divergence with a few outliers. The transition to low probability of migration is fairly rapid beyond 1% ND2 p-distance. D_XY_, an absolute measure of divergence, also scales with F_ST_ but with more outliers compared to ND2 (electronic supplementary material, figure S5). Though different measures of divergence in different DNA classes generally correlated as expected, divergence and structure of populations varied between species with no immediate patterns corresponding to geography.

### Geography and genome divergence

Relative genome divergence (F_ST_) and realized gene flow (probability of migration) is not predicted by between the current geographic definitions of allopatry and parapatry: allopatry between PNG and mainland Australia and either allopatry or parapatry within mainland Australia, respectively (electronic supplementary material, figure 2, figure S3, top; Kruskal-Wallis chi-squared=2.58e-03, df = 1, p = 0.9595). Population pairs exhibit a range of divergence levels regardless of whether the current barrier is terrestrial or marine. Divergence due to isolation-by-distance is well supported by a correlation between the adjusted F_ST_and log of the distance in km (Spearman’s rank correlation: rho = 0.427936, p = 0.004698). Correspondingly, the probability of migration also decreases with increasing distance (Spearman’s rank correlation: rho = -0.482106, p = 0.001225). Lastly, we compared the minumum landscape resistance (ie. maximum range connectivity or maximum potential for gene flow corrected for distance) between the three historic time points to F_ST_ for each population pair. All comparisons with PNG had the LGM as the period where PNG was connected with north Australia (18 pairs). The timing of maximum connectivity within mainland Australia varied: eight pairs in present-day, seven during the mid-Holocene, and nine in the LGM (electronic supplementary material, table S7, figure S3). There is a positive correlation between landscape resistance and F_ST_ (Spearman’s rank correlation test, S = 7048.6, rho = 0.4288, p = 0.004601), which translates to a negative correlation between resistance and probability of gene flow (Spearman’s rank correlation test, S = 18102, rho = -0.466829, p = 0.00183). In sum, whereas classification of allopatry and parapatry based on predictions of current distribution do not predict divergence or probability of gene flow, other factors such as geographic distance or past connectivity proves to be more appropriate predictors.

**Figure 2.**
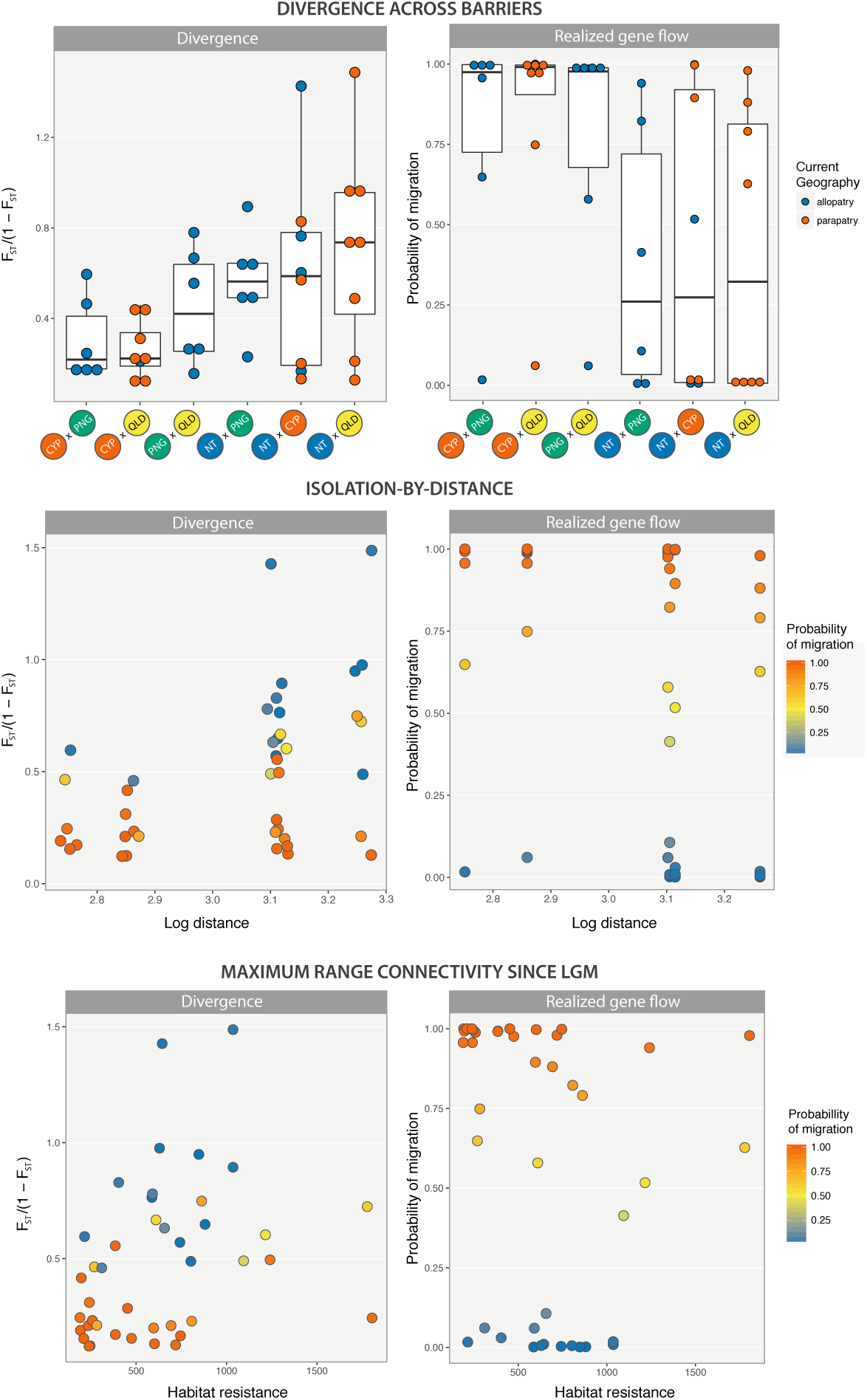
*Top*: Relative divergence through the barriers colored by whether or not population pairs have disjunct ranges. Allopatry and parapatry are defined based on connectivity in the species distribution models. The barriers are ordered based on increasing geographic distance *Middle*: Adjusted F_ST_ values plotted against log of distance to show isolation-by-distance. Color gradient depicts the relative support for the models with gene flow. Points have been jittered by 0.03 to display the number of points. *Bottom*: Divergence plotted against the highest connectivity across three time points (present, mid-Holocene, LGM).

Although the patterns described above also apply to the Z chromosome, the Z chromosome tends to have higher differentiation relative to the autosome. The difference between the autosomal F_ST_ value and Z chromosome Fst value increases as probability of gene flow decreases (Spearman rank correlation: rho = -0.5504, p = 1.58e-4) but does not correlate with current population connectivity (Kruskal-Wallis rank sum test: chi-squared = 0.12665, df = 1, p = 0.7219). This means that the amount of difference between autosomal and Z chromosome divergence is a function of overall divergence rather than spatial context. The difference between autosomal and Z chromosome values increases with increasing divergence for both relative (F_ST_) and absolute divergence (D_XY_), though absolute divergence far less so (Spearman rank correlation F_ST_: rho = 0.715672, p = 1.001e-7; Spearman rank correlation D_XY_: rho = 0.285595, p = 0.06674; electronic supplementary material, figure S6). In this system, broad sampling of population pairs reveal that difference between autosomal and Z chromosome divergence is likely simply related to level of divergence rather than population connectivity.

### Genome divergence and speciation

Designations of allopatry or parapatry in current distributions do not predict realized gene flow, even for the less diverged populations. However, our data suggests that relative divergence level influences realized gene flow throughout the entire range of divergence. We see a rapid transition from high gene flow with low divergence to low gene flow with higher divergence through a narrow range of divergence levels; similar to the model of snowballing during speciation (figure 3). Support values for the different demographic models can be found in the electronic supplementary material table S10. A subset of simulations under the strict isolation model was always recovered to have low probability of migration regardless of F_ST_ value, suggesting that the trajectory we see in our data is not likely due to artifacts in the model selection (figure 3).

**Figure 3.**
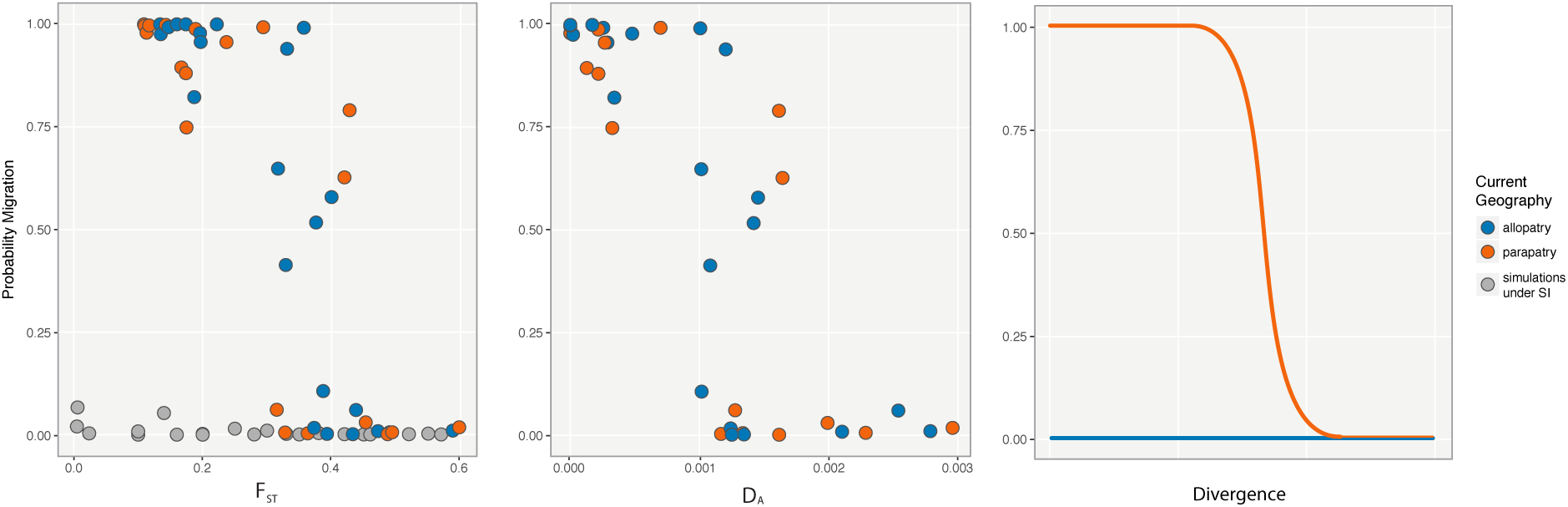
Probability of migration defined by the sum of ABC model supports for the 4 models with a migration parameter (IM, IMhetM, IMhetN, IMhetNhetM) plotted against F_ST_ and D_A_. The third panel is a plot of our expectation of the change in probability of migration under allopatry and parapatry.

After comparing our data to models proposed by Yamaguchi and Iwasa [15], all simulations retained by the ABC model selection were those simulated under the sigmoidal function representative of a snowballing effect (electronic supplementary material, figure S2). There is no support that our data follow any other proposed trajectories. The F_ST_ range corresponding to the tipping point spans ∼0.3 - 0.4. The D_A_ range corresponding to the tipping point is ∼0.1-0.17%. Species-specific points often span the entire range of the trajectory (electronic supplementary material, figure S7). Plots for the Z chromosome, D_XY_, and ND2 against probability of migration can be found in the electronic supplementary material, figure S8. Our system provides additional empirical support for a snowballing pattern in speciation.

### Genome-wide divergence

The coefficient of variance describes the distribution of the individual F_ST_ values across the RAD loci. Higher coefficient of variance corresponds to a more skewed distribution - i.e. one in which there is higher heterogeneity in levels of divergence across loci. At lower F_ST_ values (and high gene flow) there is a lower coefficient of variance as most values are close to zero. The coefficient of variance increases with increasing F_ST_ but peaks at intermediate levels of migration and starts to decrease again when the support for migration decreases. The change in the distribution of F_ST_ values follow a predictable pattern with increasing divergence where there is an initial skew from low to moderate divergence levels followed by a more uniform distribution from moderate to high divergence levels (figure 4). This may be due to similarities in genome architecture rather than differences in current geographic classification.

**Figure 4.**
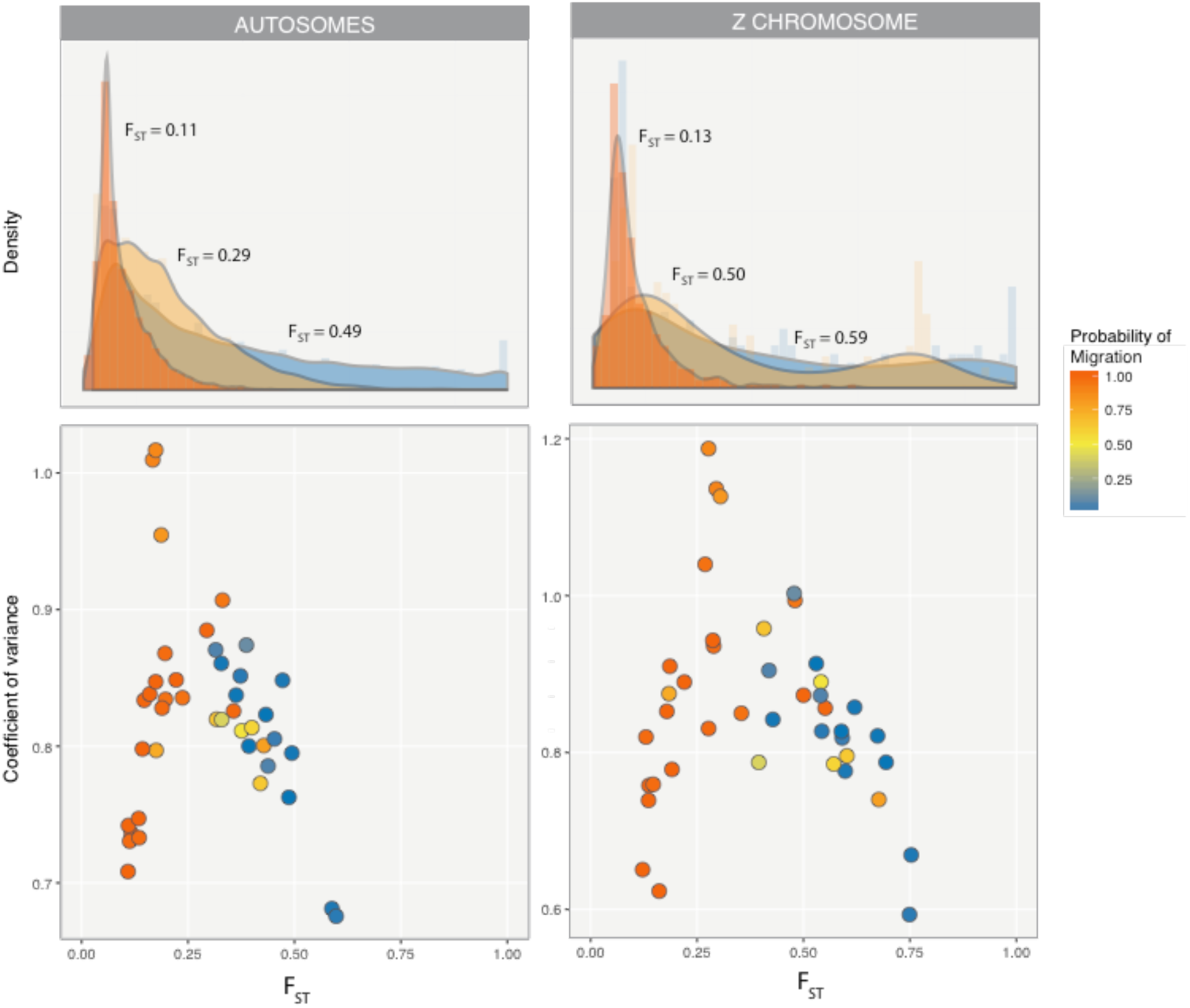
*Top*: Density of F_ST_ distributions of the ddRAD loci with increasing global F_ST_ and decreasing probability of migration. Three representative population pairs were chosen to represent different divergence levels. Low: Brown honeyeater NT & QLD, Medium: Blue-faced honeyeater CYP & QLD, high: White-throated honeyeater NT & QLD. *Bottom*: Distribution of the skew of F_ST_ distributions with increasing global F_ST_ and decreasing probability of migration.

## Discussion

Geographic mode of speciation is expected to influence the roles of gene flow, selection, and drift on divergence. However, our data suggests that although currently continuous geographic ranges should have higher potential for gene flow relative to discontinuous ranges, it does not necessarily translate to a higher probability of migration during divergence. The degree of connectivity between populations is dynamic through their evolutionary history and this results in reticulation of the genomes between those populations (figure 1, electronic supplementary material, figure S3)[59]. Our system supports the idea that current classifications of geography (allopatric, parapatric, and sympatric) are not reflective of gene flow during population divergence [6,7]. It is possible that some currently allopatric populations have high probability of gene flow reflecting past connectivity. Conversely, some currently parapatric populations may have low probability of gene flow from incompatibilities accumulated during past allopatry. Instead, there is support for geographic distance and minumum resistance being negatively correlated with realized gene flow and positively correlated with divergence. Properly classifying the geographic mode of speciation is particularly pertinent to a dynamic region like northern Australia and PNG. Despite this correlation with distance and historical connectivity, population divergence and realized gene flow of these birds across various barriers in northern Australia and Papua New Guinea span the range of the speciation continuum potentially due to lineage-specific biogeographic history or natural history (figure 3). Accumulating biogeographic studies across suture zones or shared biogeographical barriers show similar dynamics of population divergence in other avian systems as well as lizards and invertebrates [60–62].

Our data provides another empirical example in support for a snowballing pattern during speciation. There exists a certain threshold of divergence where there is a rapid transition between high and low gene flow likely resulting in speciation. Though populations at intermediate stages are still observed, they persist at a narrow range of divergence levels. Our data shows a similar rapid transition between the states of high and low likelihood for gene flow (figure 3). This transition occurs at a narrow stage of nuclear and mitochondrial divergence; F_ST_ ∼0.3-0.4, D_A_ ∼0.1-0.17%, and ND2 p-distances of ∼1-1.5%. Using the substitution rate from Pacheco et al. (2011), the ND2 p-distances would translate to 1.11 - 1.67 Mya. The range of D_A_ where transition occurs is far lower to that found in Roux et al. (∼0.5-2%) which may be a characteristic specific to avian speciation. Additional studies in other avian systems would help determine if these patterns in divergence are robust.

Unlike the nuclear loci, there are outliers in the mitochondrial data where populations with high mtDNA divergence would still have high likelihood of gene flow and low F_ST_ (electronic supplementry material, figure S4). These populations are from the dusky myzomela (CYP & QLD, CYP & PNG, and PNG & QLD) and from the white-throated honeyeater (CYP & QLD). The dusky myzomela population pairs are part of the small subset which are all allopatric on mainland Australia. One possible explanation for the white-throated honeyeater is that gene flow after secondary contact has homogenized the nuclear genome but maintained the local ND2 haplotypes in accordance with Haldane’s rule [63].

Although neither ecological selection nor incompatibility loci were explicitly tested, this sigmoidal trajectory of speciation is consistent with the parapatric models (divergence-with-gene flow) proposed by Yamaguchi and Iwasa [15]. As suggested by the dynamic geographic history, population pairs likely experienced varying rates of gene flow through time [8,59]. On the other hand, it has also been shown that a nonlinear accumulation of divergence can occur under certain neutral scenarios [20]. Ideally, estimates of population splitting time would inform us about the timing and duration of speciation and therefore the rate of accumulating divergence; however, we found that we could not use ddRAD to infer divergence times reliably [64].

Within the genome, there is variation of divergence across loci owing to various degrees of standing genetic variation and recombination [25–27]. The change in the distribution of F_ST_ values across loci is a coarse estimate of the change in landscape of divergence through time.

Although the L-shape or the skew of F_ST_ distributions between parapatrically diverging populations have been attributed to a few loci under divergent selection with the rest homogenized by gene flow [30,31], our broader sampling of population pairs show that the change in F_ST_ distribution follow a predictable pattern with increasing F_ST_ or decreasing probability of gene flow regardless of connectivity. It is also likely that this pattern of accumulation of divergence is due to variation in nucleotide diversity across the loci [20,25]. The change in skew with increasing F_ST_ could also be driven by linked selection instead of resistance to gene flow as recent studies have shown [27,65]. It would be important to complement this study with detailed studies across hybrid or suture zones to disentangle the role of gene flow and linked selection on divergence between hybridizing populations.

Finally, we note some important points for the study of the geographic mode of speciation. First, neither range overlap nor migration rate is static during speciation and it is more likely that populations experienced various degrees of geographic connectivity and gene flow through time [59]. This is particularly pertinent for highly vagile taxa, like birds, in highly dynamic geographic regions, like the Australopapuan region. Additionally, populations that are currently allopatric due to vicariance could have experienced periods of reduced but ongoing gene flow during the formation of the barrier [66]. Second, population differentiation would also vary if a population has accumulated isolating mechanisms during allopatry and accelerated divergence during secondary contact (alloparapatry) [2]. Depending on the particular demographic history, isolation with continuous migration (primary divergence) can be difficult to distinguish from gene flow after secondary contact from genetic data alone [24]. Third, population divergence in allopatry may not necessarily translate to speciation. Seeing that most species definitions rely on degree of reproductive isolation there are no consistent criteria to differentiate discontinuous populations with no gene flow from allopatric species [1,67,68].The combination of reduced gene flow, either completely or partially, and genetic drift may result in population divergence but other factors likely play a more important role in speciation.

## Conclusion

Our comparative study of divergence of bird populations through the speciation process highlights the dynamics of geographic history and their influence on divergence. Further, it provides additional support for a snowballing pattern in speciation, and it characterizes broad patterns of genomic divergence through time. The divergence that we discuss in this paper is presumed to be neutral and therefore would benefit from a replicate study looking at divergence in coding regions to observe whether different marker types have different trajectories in the same populations. The results of this broad study also clearly describe the pattern of accumulation of divergence which lend support to the emerging relevance of linked selection in genome divergence. Comparative studies, particularly with multiple species in a shared geographic region, help elucidate patterns of genome divergence during speciation. To fully comprehend the patterns of neutral and adaptive genomic divergence, we need to sample broadly both phylogenetically and geographically thus affecting shared patterns in speciation differentiated from the exceptions.

## Author contributions

JVP conceived ideas and designed the project, collected the data, performed the analyses, and led the writing of the manuscript. LJ and CM provided ideas during the analysis and writing of the manuscript

### Acknowledgments

We would like to thank Ian J. Mason and Alex Drew for greatly appreciated help with collecting samples in the field, Alexander Xue for assistance with demographic analyses, Matteo Fumigalli for assistance with ngsTools, and Daniel Rosauer for assistance with species distribution modelling. We would also like to thank Sonal Singhal, Sally Potter, and Daniel R. Wait for helpful discussions and comments on the manuscript. Lastly all data collection was carried out in Australian National University’s Biomolecular Research Facility and most analyses were carried out in the ABC Bioinformatics Development Cluster.

